# Screening human embryos for polygenic traits has limited utility

**DOI:** 10.1101/626846

**Authors:** Ehud Karavani, Or Zuk, Danny Zeevi, Gil Atzmon, Nir Barzilai, Nikos C. Stefanis, Alex Hatzimanolis, Nikolaos Smyrnis, Dimitrios Avramopoulos, Leonid Kruglyak, Max Lam, Todd Lencz, Shai Carmi

## Abstract

Genome-wide association studies have led to the development of polygenic score (PS) predictors that explain increasing proportions of the variance in human complex traits. In parallel, progress in preimplantation genetic testing now allows genome-wide genotyping of embryos generated via *in vitro* fertilization (IVF). Jointly, these developments suggest the possibility of screening embryos for polygenic traits such as height or cognitive function. There are clear ethical, legal, and societal concerns regarding such a procedure, but these cannot be properly discussed in the absence of data on the expected outcomes of screening. Here, we use theory, simulations, and real data to evaluate the potential gain of PS-based embryo selection, defined as the expected difference in trait value between the top-scoring embryo and an average, unselected embryo. We observe that the gain increases very slowly with the number of embryos, but more rapidly with increased variance explained by the PS. Given currently available polygenic predictors and typical IVF yields, the average gain due to selection would be ≈2.5cm if selecting for height, and ≈2.5 IQ (intelligence quotient) points if selecting for cognitive function. These mean values are accompanied by wide confidence intervals; in real data drawn from nuclear families with up to 20 offspring each, we observe that the offspring with the highest PS for height was the tallest only in 25% of the families. We discuss prospects and limitations of PS-based embryo selection for the foreseeable future.

## Introduction

The use of biotechnology to influence the genetic composition of human embryos in the absence of specific disease risk raises many ethical concerns, and the recent live births resulting from human embryonic CRISPR editing has heightened global attention to these issues [1,2]. Currently, the most practical approach to genetic “enhancement” of embryos is preimplantation genetic screening of IVF embryos. Preimplantation genetic diagnosis and screening [3] have been utilized for years to avoid implantation of embryos harboring monogenic disease-causing alleles or aneuploidies. Recently, it also became technically feasible to generate accurate genome-wide genotypes from single-cell input [4]. This development, coupled to recent progress in complex traits genetics, made it possible to genetically screen embryos for polygenic traits, and has raised the prospect of “designer babies” [5].

Perhaps the most controversial potential application of polygenic embryo selection would be selection for intelligence, especially given the abhorrent history of the early-20^th^ century eugenics movement [6]. While most ethicists are deeply troubled by such prospects, at least one prominent scholar has suggested that there is an ethical obligation for parents to “select the best children” [7]. In our view, any discussion of the ethics of embryo selection must be informed by quantification of the expected utility of polygenic selection, either as of today, or as reasonably projected into the future. In this report, we thus utilize statistical and empirical methods to evaluate the potential effects of human embryo selection for polygenic traits.

Polygenic scores (PS) are derived from large-scale genome-wide association studies (GWAS) of complex traits, which can be quantitative (such as intelligence or height) or categorical (such as disease status, in which case they are often referred to as ‘polygenic risk scores’) [8]. A PS is the count of effect alleles in an individual’s genome, weighted by each allele’s strength of association with the trait of interest in an independent GWAS [9]. The predictive power of a PS is usually represented by 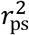, or the proportion of variance of the quantitative trait explained by the PS. To date, the largest GWAS of intelligence [10,11] has demonstrated a relatively modest out-of-sample 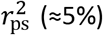, despite large sample sizes (*n*≈300,000 individuals). By contrast, recent large-scale GWASs of height have attained 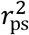 of approximately 25%, while demonstrating a highly polygenic genetic architecture similar to intelligence [12]. Consequently, in the present report, we analyze height in addition to cognitive function, which also allows us to exploit several datasets in which height data, but not intelligence data, are available.

PSs are typically evaluated on a cohort basis, and are not used to differentiate one individual from another (although a recent report has demonstrated that, for an extraordinarily tall NBA player, the PS for height was >4 standard deviations above the population mean [13]). In order for polygenic embryo selection to hold potential utility (independent of ethical considerations), PSs must provide sufficient predictive power to differentiate between embryos within the restricted range of genetic variance available in a single family, and with a finite number of embryos. Two reports utilizing only mathematical modeling have suggested that substantial effect sizes for embryonic selection are possible [14,15]. But to our knowledge, despite the widespread application of polygenic scores to complex traits and precision medicine in the research literature [16], no published studies have empirically examined the possibilities and limitations of a polygenic approach to embryo selection.

We consider here embryo selection in the context of a hypothetical IVF cycle. Our quantity of interest is the difference between the predicted value of the selected trait (i.e., height or intelligence) when the embryo with the highest PS is selected, compared with the value of the average embryo (i.e., the mean across embryos). We term this difference the *gain*, and we further differentiate between the *predicted* gain, as determined by the PS, and the *realized* gain, as observed in the fully-grown offspring. Because no study can be performed in actual embryos, we utilize three sources of data: 1) a quantitative genetic model; 2) simulated embryo genomes generated using realistic parameters from existing genotyped datasets of adults with known phenotypic values; and 3) a unique pedigree dataset of nuclear families with large numbers of offspring (10 on average), now fully-grown adults, with available genotype and phenotype data. In our simulated data, we examine the gain as a function of varying predictive strengths 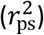 of the PS, as well as of the number of embryos (*n*) available; these results were compared against a theoretical model derived for average gain. Although a typical IVF cycle may produce 3-8 viable embryos (median=5; [17]), we examine the gain across a broad range of values of *n*, given the possibility of future advances in IVF technology. Particular emphasis is placed on *n* = 10, representing a plausible upper bound within the foreseeable future.

## Results

We first developed a simple quantitative genetic model for the expected gain. The model assumes a polygenic additive trait with no assortative mating, and hence no correlation between the scores of SNPs between homologous chromosomes or chromosomes of spouses. We recognize that statistically significant assortative mating has been demonstrated for genetic variants associated with polygenic traits such as height and educational attainment [18]; however, the overall magnitude of this effect accounts for <5% of the variance in spousal phenotype [19,20]. Assortative mating would tend to reduce the efficacy of embryo selection due to reduced variance available from which to select, and thus our results described below represent an upper bound on the potential gain.

We assumed a couple has generated *n* embryos, and computed the distribution of the polygenic scores of the *n* embryos for a trait with phenotypic variance 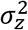, of which a proportion 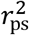 is explained by the PS. The set of *n* polygenic scores can be modeled as having a multivariate zero mean normal distribution with all variances equal to 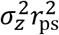 and all covariances equal to 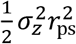. The *gain* is formally defined as the difference between the maximal and average PSs among the *n* embryos. Based on properties of multivariate normal distributions, the mean gain can be shown to be approximately (for details see the **Supplementary Note**)

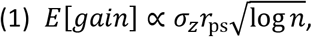

where the coefficient of proportion is ≈0.77. A more accurate formula based on extreme value theory can also be derived (**Supplementary Note** Eq. (35)). Most notably for our purposes, the mean gain increases with the square root of the variance explained (or linearly with the correlation coefficient between the PS and the trait), but the effect of *n* is considerably attenuated, as denoted by the square root and log transformation in Eq. (1).

Next, for our simulations, we used genotypic and phenotypic data from two cohorts. The Longevity cohort contained 102 couples of Ashkenazi Jewish origin with genome-wide genotypes and information on height, drawn from a larger longevity study [21]. The ASPIS cohort [22] contained 919 young Greek males with genome-wide genotypes and information on general cognitive function. To simulate embryos, we used either actual couples (for the Longevity cohort) or randomly matched couples (for both cohorts), and generated *n* = 10 or 50 synthetic offspring per couple based on a standard model of recombination (see *Methods* for details).

To predict the height or IQ of each embryo, we used polygenic scores based on summary statistics derived from recent large-scale GWAS meta-analysis. For height, the most recent meta-analysis contained ≈700,000 individuals [12] and did not include the subjects in our test (Longevity) cohort. For IQ, we utilized the most recent published meta-analysis [11], from which the COGENT set of cohorts (including the ASPIS cohort) had been removed, resulting in a discovery sample size of *n* = 234,569. We optimized the polygenic scores with respect to imputation, LD-pruning, and the P-value threshold (*Methods*). Our scores predicted height in the Longevity cohort with 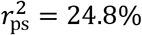 and IQ in the ASPIS cohort with 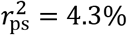, both within one percentage point of the maximum out-of-sample predictive power reported in the original GWAS. Using linear regression of the phenotype (age- and sex-corrected for height) on the polygenic scores in each cohort, we predicted the height or IQ of each simulated embryo.

Having calculated the predicted height of each simulated embryo from the Longevity cohort and the predicted IQ of each simulated embryo from the ASPIS cohort, we sought to test the predictions of the mathematical model in Eq. (1). To examine the relationship between predicted gain and the variance accounted for by the PS, we fixed the number of embryos to *n* = 10, and plotted the mean gain for height against increasing 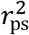. Because polygenic contributions to most complex traits (including height and IQ) are evenly distributed throughout the genome [23], we generated PSs that were progressively stronger using PSs derived from growing subsets of the 22 autosomes (e.g., chromosome 1 SNPs only, chromosome 1 + chromosome 2 SNPs only, etc.). As shown in Figure 1, the average gain reaches ≈3cm or ≈3 IQ points when the full genome-wide PS is used (corresponding to ≈0.5 and ≈0.2 standard deviations of the trait, respectively). The average gains obtained from varying 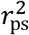 are close to the values predicted by the theoretical model (Eq. (1)). Our results did not differ when the actual couples are used as the source of the simulated embryos (Figure 1, center), compared to couples randomly matched from the Longevity cohort (Figure 1, left), indicating that effects of any assortative mating in this dataset are *de minimis*.

**Figure 1.**
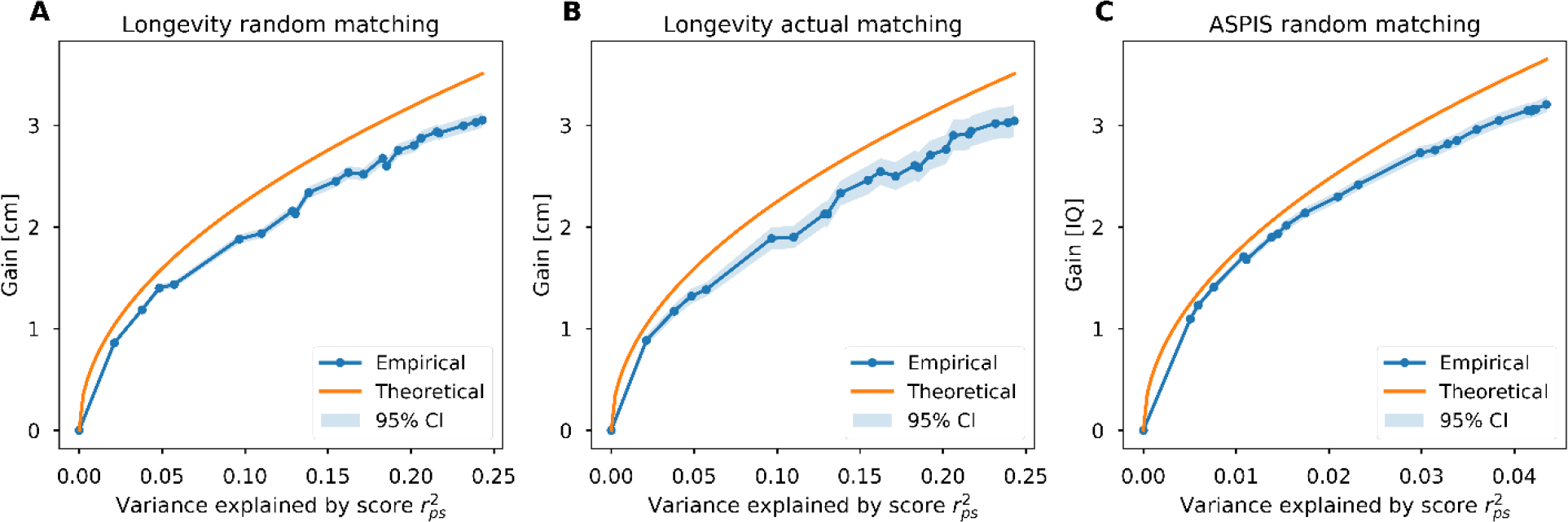
The mean gain vs the proportion of the variance explained by the PS. Blue dots and the 95% confidence intervals represent simulations with 10 embryos per couple. To generate scores with increasing proportions of variance explained, we gradually added chromosomes 1 to 22 to the computed PS. The orange line corresponds to the theoretical model derived in the **Supplementary Note** and described in Eq. (1). The 95% confidence interval, for each value of 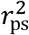, is based on ±1.96 the standard error of the mean over the simulated families. **(A)** Gain in height for random couples: 500 simulated pairings drawn from the Longevity cohort. **(B)** Gain in height for actual couples: 102 couples from the Longevity cohort. **(C)** Gain in IQ for random couples: 500 simulated pairings drawn from the ASPIS cohort. Results were averaged across couples in all panels.

The PSs used so far are based on current GWAS results and on a simple LD-pruning and P-value-thresholding strategy. However, GWASs are expected to increase in size (in particular given the rapid growth of the direct to consumer genetic industry [24]), and statistical prediction methods are constantly improving [e.g., [25–28]]. Given that the theoretically predicted relationship of gain with *r*_p*s*_ was supported by the data in Figure 1, we can forecast the prospects of embryo selection as predictors become increasingly accurate. For example, doubling the proportion of explained variance of height from ≈25% to 50% is expected to increase the mean gain from ≈3 to ≈4.24cm, with a maximum possible gain of ≈5.5cm for 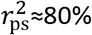 (the upper bound of the heritability of the trait, as derived from twin studies; [29]). Similarly, quadrupling the variance explained for IQ would lead to a doubling of the gain, to ≈6 IQ points (given *n* = 10 embryos).

Next, we tested the relationship between the gain and the number of embryos, holding 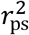 constant. In Figure 2, we show the expected gain vs the number of embryos, for up to 50 embryos. Comparison to the theoretical model again shows good agreement, with an even better fit demonstrated in Supplementary Figure 1 based on a more accurate approximation (**Supplementary Note** Eq. (35)). Two implications are immediately apparent from Figure 2. First, current reproductive technologies are in the most sensitive area of the curve. With a typical IVF cycle yielding 5 testable, viable embryos [17], the predicted gain is reduced from ≈3 to ≈2.5 (cm or IQ points); below 5 embryos, the gain drops precipitously. Second, there is a rather slow increase of the mean gain as the number of embryos increases beyond 10. Thus, even with 1000 embryos, the mean gain would be only ≈1.7 times higher compared to selection with 10 embryos. Again, no differences were observed between randomly paired and actually married couples (panels A and B). The pattern for intelligence was roughly equivalent to that observed for height (panel C).

**Figure 2.**
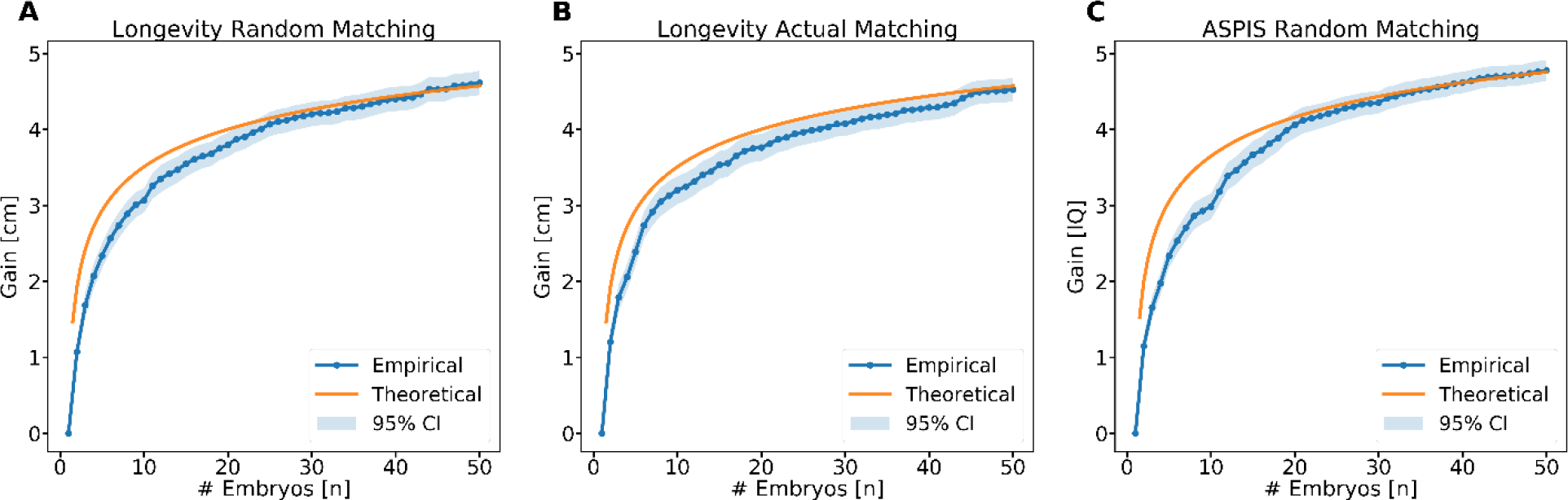
The mean gain vs the number of embryos. Blue dots are from simulations, and orange lines are for the theoretical prediction (Eq. (1)). All details are as in Figure 1.

Both of the results above demonstrate the *average* gain expected under varying levels of 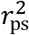 and *n* across 102 real couples or 500 simulated couples. However, for any given couple, the predicted gain will further vary around this mean. The distribution of the gain, when choosing the best out of 10 embryos, is shown in Figure 3 for height (for both random and actual couples) and IQ. The gain in height is typically between 1-6cm, with a median of 2.88cm for random couples (IQR: 2.34-3.80) and 3.02cm (IQR: 2.43-3.84) for actual couples. The gain in IQ was between 1-7 points (IQR: 2.43 - 3.84), with a median of 3.02 IQ points. Thus, the predicted gain for a given couple may be somewhat higher or lower than suggested by the mean results of our simulations, due to variation across couples and the random assortment of SNPs in the offspring.

**Figure 3.**
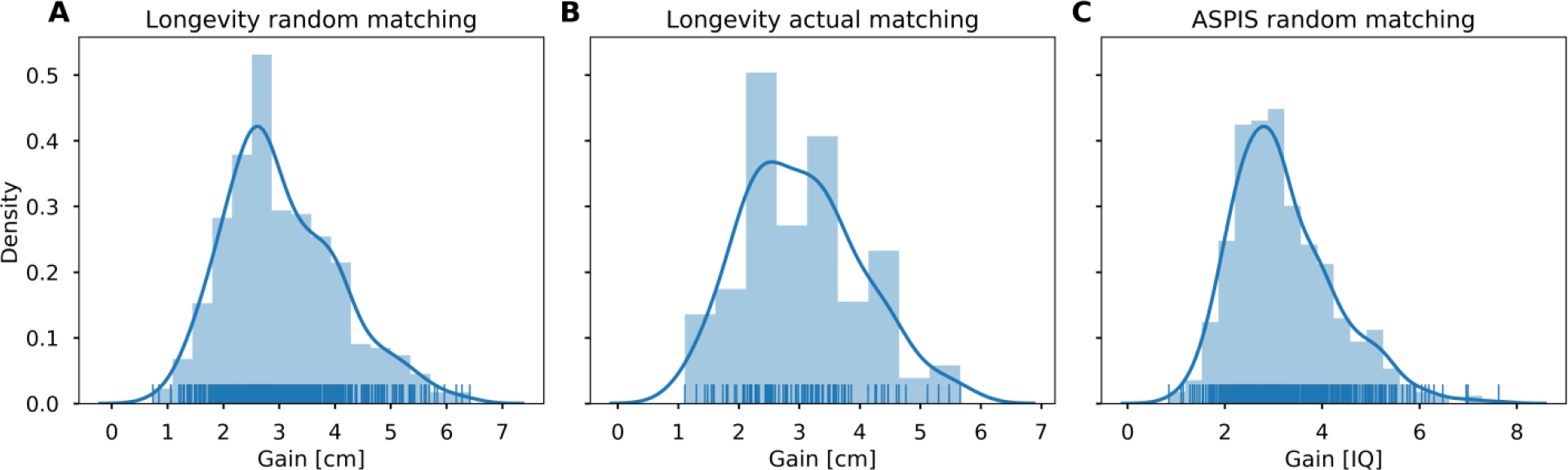
The distribution of the predicted gain from embryo selection with 10 embryos per couple. **(A)** The gain in height by simulating 500 random couples from the Longevity cohort. **(B)** Same as (A), but with actual spouses (*n* = 102). **(C)** The gain in IQ by simulating 500 random couples from the ASPIS cohort. Lines are estimated densities.

The variance depicted in Figure 3 represents the variability of the *predicted* gain across couples, but environmental variance leads to additional and substantial variability in the *realized* gain, as observed in the phenotype of the offspring. A simple calculation (**Supplementary Note**, Section 4.2) shows that given a predicted gain, the 95% prediction interval for the (zero-centered) trait value is approximately

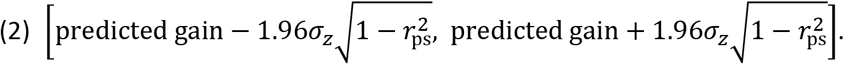

Eq. (2) can be compared to a 95% prediction interval of [−1.96*σ*_*z*_, 1.96*σ*_*z*_] without selection. Currently available PSs account for substantially less variance in phenotypic values than expected by the heritability, in part due to rare genetic variation not captured by current GWAS [30]. However, prediction intervals can be narrowed based on the parental phenotypic values, which are usually known. For example, it has long been known that mid-parental height can explain ≈40% of the variance in height of the offspring [31], or theoretically *h*^4^/2 ≈ 32% [32]. However, these ≈32% of the variance overlap with the ≈25% explained by the PS, and the combination of both sources of information can never explain more than the heritability. As shown in Figure 4A, even under the extreme scenario where the *combination* of the PS and the parental values explain the entire heritability of height (≈80%), there would still be ±5cm interval around any predicted gain due to environmental and stochastic factors. Based on either the current PS alone, or based on the parents alone, the interval would be as large as ±9-10cm. For IQ, the 95% prediction interval would be ±13-19 points in case the entire heritability is explained (assuming *h*^2^ ∈ [0.6,0.8]), or ±24-27 points based on the parents (Figure 4B). Thus, the unexplained variance yields a wide confidence interval around any predicted value for an offspring’s height, and therefore a considerable uncertainty in the realized gain that any given couple can expect from embryo selection. This would need to be combined with the variability in the predicted gain itself, as depicted in Figure 3, thereby substantially attenuating any guarantees for the potential benefit.

**Figure 4.**
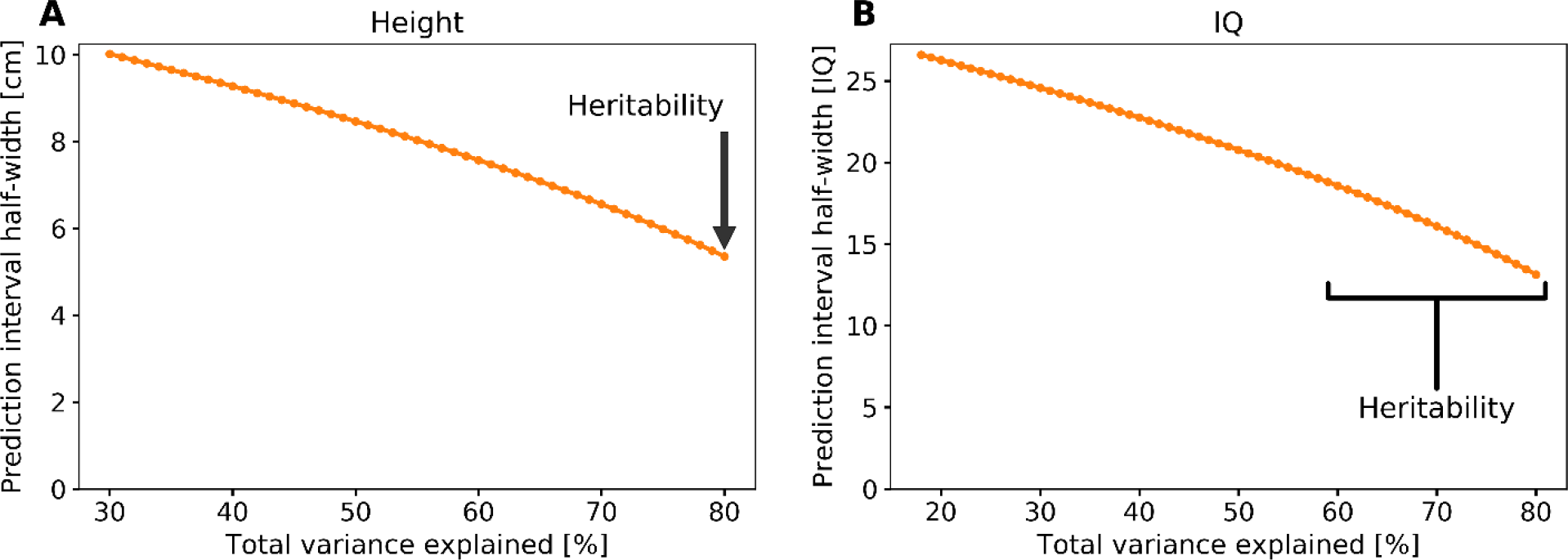
The prediction interval width as a function of the proportion of variance explained by the combination of parental phenotypes and the PS of the child. If the proportion of variance explained is *p*, the half-interval width is 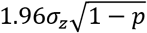. **(A)** The prediction interval for height, assuming *σ*_*z*_ = 6cm. The proportion *p* is unknown, but cannot exceed the heritability, which we assume to be *h*^2^ ≈ 0.8, and cannot fall under *h*^4^/2 ≈ 0.32, which is the theoretical variance explained by the mid-parental height. **(B)** The prediction interval for IQ, assuming *σ*_*z*_ = 15 points. We assume the heritability is in the range [0.6,0.8], with a minimal variance explained of 0.6^2^/2 = 0.18.

To demonstrate the implications of the above equations, consider the extreme case in which the variance explained by the PS is so large that the contribution from the parents’ phenotypes is negligible and Eq. (2) is applicable, with the predicted gain further set to its mean value. For height, with 70% of the variance explained and selecting out of 10 embryos, a 95% prediction interval for the height of a male child (assuming 175cm for the population average, an SD of 6cm, and a normal distribution) would be approximately 180±6cm (i.e., 174-186cm). This is compared to 175±12cm (163-187cm) without selection. For IQ (mean 100 and SD 15), with 30% of the variance explained, the 95% prediction interval would be approximately 109±25 (84-134), compared to 100±30 (70-130) without selection. Even under this extreme case, the future child has a non-negligible probability (≈0.26, assuming a normal distribution) to have an IQ below the population average.

Finally, to evaluate the utility of embryo selection in a real-world setting, we examined PS for height in a unique cohort of 28 large families with up to 20 offspring each (range 3-20; mean=9.6), now grown to adulthood. While all these families were the result of traditional means of procreation, we treated the offspring data as if all offspring were simultaneously generated embryos available for selection based on their PSs. Figure 5A depicts the actual difference in height between the offspring with the highest PS, compared to the average height of all the offspring in each family, i.e., the *realized* gain. (All heights were corrected for age and sex). While the observed values average around the mean gain predicted by the theory, there was substantial variability in the realized gain. Some families realized a gain of up to 10cm, while for 5 of the 28 families, choosing the embryo with the highest PS would have resulted in an offspring with height below the average (i.e., gain < 0).

**Figure 5.**
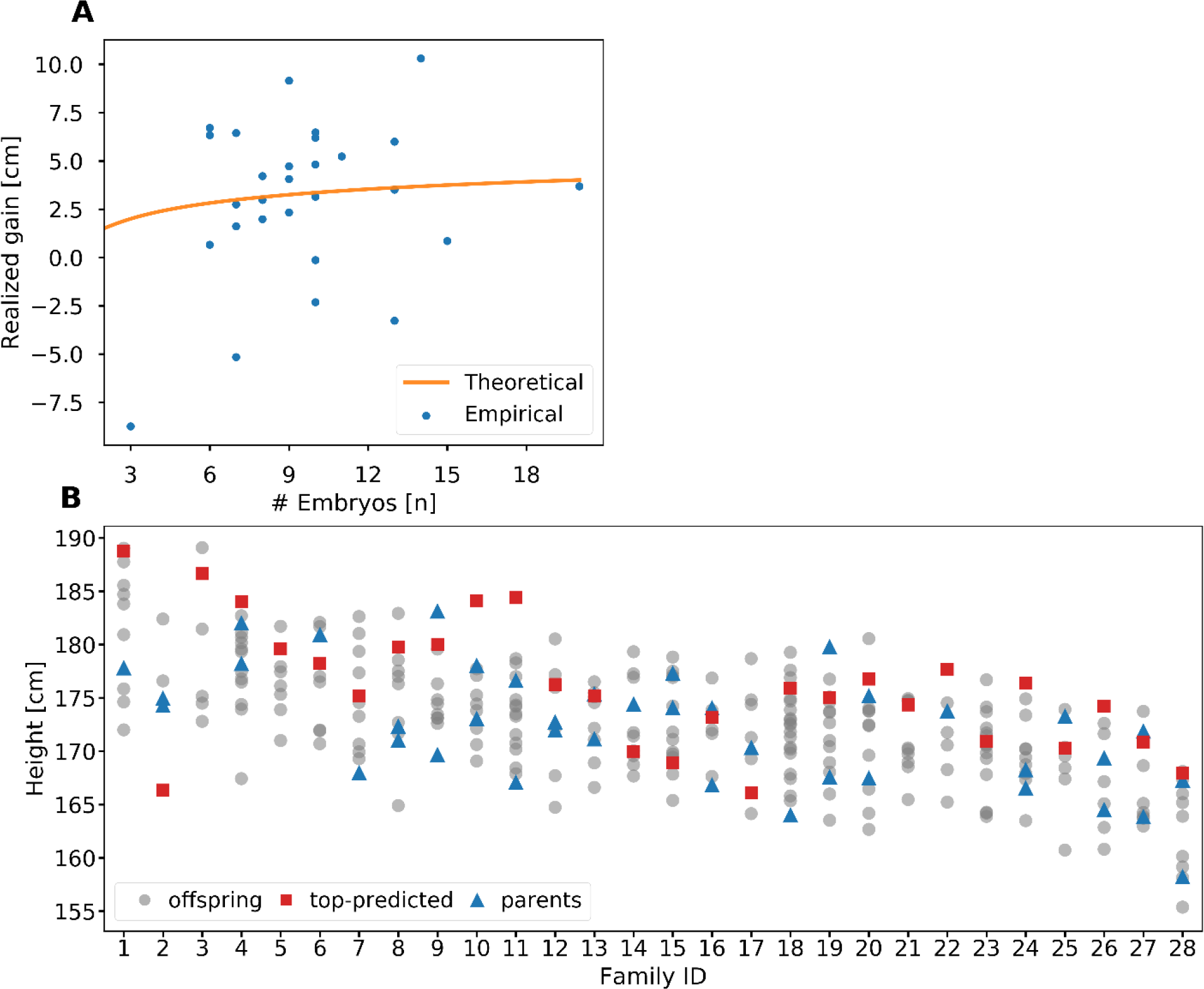
Analysis of selection for height in 28 real families with up to 20 adult offspring each. **(A)** The realized gain in each family, defined as the difference between the actual (age- and sex-corrected) height of the offspring with the highest PS and the average height of all offspring in the family. The theoretical prediction is based on Eq. (1). **(B)** The actual height (age- and sex-corrected) of all members of all families. The figure demonstrates the effect of the current low-accuracy prediction models, as the tallest-predicted sibling (red squares) is usually not the actual-tallest sibling (only 7/28 times). Siblings are depicted as grey dots, and the parents of each family as blue triangles. In some families only one parent was available.

The inherent uncertainty in PS-based selection is also demonstrated in Figure 5B, which displays the actual height for each family member. It is notable that the offspring with the highest PS (red squares) is the tallest actual offspring in only 7 of the 28 families. Moreover, when repeatedly downsampled to *n* = 7 children, the offspring with the highest PS was the tallest in ≈31.5% of the families, close to the theoretical prediction (≈33.4%; **Supplementary Note** Section 4.4). Across all families, the tallest child was on average ≈3.0cm taller than the child with the tallest predicted height, again very close to the theoretical prediction (3.1cm; **Supplementary Note** Section 4.3).

## Discussion

In this paper, we explored the expected gain in trait value due to selection of human embryos for height and IQ. We showed that the average gain, with current predictors and with 5 viable embryos, is around ≈2.5cm and ≈2.5 IQ points. We predicted and confirmed by simulations that the gain will increase linearly with the square root of the variance explained by the predictor, but much more slowly with the number of embryos. These results contrast with the only two studies addressing this question to date, both of which employed only mathematical modeling; those studies suggested that much larger effect sizes were possible with currently available scores and technologies [14], and that increasing the number of available embryos would have the largest effect on potential gain [15]. The only empirical study comparable to this report was an examination of PS in the prediction of milk yield in dairy cattle [33]. In 17 sets of approximately ≈6 tested embryos, the top scoring embryo had an expected gain of approximately 5% of the trait value (≈0.35 standard deviations) compared to the average embryo. Since the currently available PS for milk yield has comparable 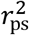 to that for human height, it is reassuring that the reported gain is similar to that reported here.

Given that 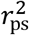 holds the strongest effect on potential gain from embryo selection, it is worthwhile to consider the potential for increasing 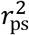 for height and IQ in the foreseeable future. Increasing sample sizes of discovery GWASs is the most straightforward means of increasing 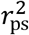 [34]. For educational attainment, a trait strongly correlated with IQ (*r*_*g*_ ≈ 0.70; [35]), increasing GWAS sample size from ≈300K [36] to ≈1.1M [37] resulted in a proportional increase in out-of-sample variance explained, from 3.2% to 11%. However, the variance explained by the predictor is not expected to increase linearly with the GWAS sample size [38]. For height, the maximum out-of-sample 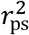 only increased from 17% to 24.6%, despite a near-tripling of discovery GWAS sample size from ≈250K individuals [39] to ≈700K individuals [12].

Second, 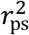 can be enhanced by the addition of increasingly rare variation to the discovery GWAS [30], especially since negative selection results in larger per-allele effect sizes at the lower end of the frequency spectrum [40]. Current imputation panels are limited in their ability to accurately assess variants with frequencies below 1%, but will continuously improve as imputation panels increase in size and representation of varying populations [41,42]. For example, a recent family-based study [43] has demonstrated that more than half of the variation in cognitive ability is attributable to rare variation not captured by current GWASs (see also [44]). Importantly, very little of this variation is private to individual families; most could be captured by population-based reference panels of sufficient size to accurately impute variants at 0.1% minor allele frequency and greater.

Third, statistical approaches to calculating PSs from GWASs are becoming increasingly sophisticated [16,45]. Most notably, the application of penalized regression methods to the generation of PSs holds the potential for rapid gains in 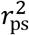 without requiring any additional data collection in either GWAS datasets or imputation reference panels [26,46]. For example, initial evidence suggests that currently available datasets might be able to explain up to 50% of the variance in height by using LASSO, and that a similar doubling of explained variance is also possible for cognitive phenotypes [27]. Additionally, the use of multiple related phenotypes has been demonstrated to enhance the predictive power of PS [47]; for example, the combination of educational attainment and intelligence GWAS may permit a doubling of cognitive 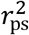 [48]. Finally, it has recently been suggested that enrichment of certain subcategories of functional variation (e.g., coding, conserved, regulatory, and LD-related genomic annotations) in GWAS results can be leveraged to further enhance prediction accuracy [49,50].

While it is likely that some combination of the above factors will increase the accuracy of PSs in the near future, substantial limitations to PSs must also be acknowledged [51]. First, PSs do not account for extremely rare Mendelian variants associated with extreme phenotypes such as short stature [52] or intellectual disability [53]. More broadly, the lower end of the phenotypic distribution is less well predicted from common variant PS than the middle and upper percentiles [54]; this fact limits the utility of PSs for “reverse” embryonic selection (i.e., to avoid extreme low values). Second, it is well known that PSs lose substantial power, or may even be invalid, when applied across different populations [55–57]. Moreover, even within a single population, subtle remaining ethnic and geographic stratification effects may result in inflated estimates of 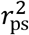 [58–60], limiting applicability to individual prediction. Third, SNP effects may be environmentally sensitive, and may not be consistent across time and place [61].

Beyond these limitations in PS power and accuracy, several additional constraints on the expected utility of embryo selection are notable. First, we did not explicitly model assortative mating, which likely exists to some extent for traits such as height and cognitive ability [18,62], and is expected to reduce the potential available variance for embryo selection. While there was no detectable effect of assortative mating in our Longevity cohort, these subjects represented an older birth cohort, and assortative mating on phenotypic traits may be increasing. Second, the number of embryos per IVF cycle is usually less than 10 [17], and, as can be seen in Figure 2, in this regime the utility drops sharply with a decreasing number of embryos. Third, with the increasing age of childbearing, so does the increase in the proportion of aneuploid embryos. For example, the proportion of aneuploid embryos is 35% for women aged 35 and 60% at age 40 [63]. Relatedly, embryos with particularly high polygenic scores are not guaranteed to implant and lead to a live birth. While it is theoretically possible to perform multiple IVF cycles to generate more embryos, IVF is invasive, involves a substantial discomfort to the prospective mother, and requires significant financial means [64] (which would often also mean an older age of the prospective parents and fewer viable embryos per cycle). To the best of our knowledge, no upcoming technology is expected to significantly increase the number of oocytes extracted per IVF cycle [65,66]. While it has been suggested that induced pluripotent stem cells may greatly increase the potential number of available embryos [67,68], such technologies are not close to implementation for human reproduction. Either way, even with tens of viable embryos, our simulations show that the gain in trait value would be relatively small (Figure 2).

Perhaps more importantly, we have demonstrated that two sources of variability result in wide confidence intervals for the prediction of final observed phenotypic values: 1) the random assortment of SNPs will result in variability of the predicted gain around its mean value; and 2) environmental variation will produce considerable additional uncertainty around the predicted gain. In our empirical dataset, the majority of offspring who were the tallest among their siblings were not those with the highest PS, and a substantial fraction of “selected” offspring had lower than average phenotypic values. Regardless of the future accuracy of 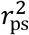 or the number of available embryos, these uncontrollable sources of variability will limit the appeal of selection for any individual couple.

A final reason for caution over the utility of embryo selection is the widespread pleiotropy across most traits [69–71]. For example, IQ is negatively correlated with most psychiatric disorders [72], but is positively correlated with autism and anorexia [73]. Therefore, selecting an embryo on the basis of higher predicted IQ will increase the risk for autism or anorexia in the offspring. In general, once IVF and genotyping/sequencing have been performed, couples may desire to attempt to select for multiple phenotypes, as well as for a reduced risk for various diseases. This will in turn lead to smaller gains per each individual trait.

Finally, we note that in this paper we did not consider the prospects, nor the ethics, of “population-scale” embryo selection for IQ or other traits. While claims were made that population-scale selection could lead to a dramatic increase in trait values at the population level [74], we leave a rigorous evaluation of this prediction to future studies. Additionally, we do not consider here the ethical, moral, and legal underpinnings and consequences of embryo selection [75,76]. We hope that this work will promote an open and evidence-based debate of these aspects among the public and policymakers.

## Methods

### Cohorts for simulating offspring

#### Longevity

Our data included 208 individuals from 104 couples who were part of the LonGenity study of longevity and aging in Ashkenazi Jews (the “Longevity” cohort). Genotyping was performed using Illumina HumanOmniExpress array. Genotyping and QC were previously described [77–80]. The number of SNPs was 704,759, with an average missing rate 0.2%. We removed duplicate variants and variants with missing rate >1%. Height was available for all individuals except two who were discarded along with their spouses. Height was 177±6cm (mean±SD) in males (range 163-191) and 163±6cm in females (range 147-175).

#### ASPIS

The Athens Study of Psychosis Proneness and Incidence of Schizophrenia [22] (henceforth “ASPIS”) included 1066 randomly selected young male conscripts aged 18 to 24 years from the Greek Air Force in their first two weeks of admission. All participants were free of serious medical conditions. Cognitive measures included: Raven Progressive Matrices Test (Raven Matrices; raw score); Continuous Performance Task, Identical Pairs version (CPT-IP; d-prime score); Verbal N-Back working memory task (Verbal NBack; total accuracy); and Spatial N-Back working memory task (Spatial NBack; total accuracy). General cognitive ability scores were generated using the first principal component from a Principal Components Analysis. Genotyping was performed on Affymetrix 6.0 arrays [81–83]. The number of SNPs was 487,126, with an average missing rate 0.3%. Out of the 1066 genotyped samples, 147 had their cognitive function scores missing and were discarded from the analysis, leaving 919 individuals. We transformed the scores to IQ points by scaling the mean to 100 and the standard deviation to 15 (range 47-140).

#### Phasing

We phased both cohorts (separately) using SHAPEIT2 [84]. Default parameters were used, except for using 200 states (to improve precision), and an effective population size of 12k, similar to the value suggested for Europeans. The genetic map used was from HapMap [85].

### Polygenic score (PS) calculation and phenotype prediction

#### Height

We used summary statistics from [12], a meta-analysis based on [39] and the UK Biobank [86]. Effect sizes were available for 2,334,001 SNPs, of which 1,789,210 were missing from the Longevity panel. Another 241 variants had mismatching alleles, leaving a total of 544,550 for downstream analyses. Scoring of individuals based on the summary statistics was performed in PLINK [87] with the no-mean-imputation flag.

Given a PS, we predicted height in a two-step approach. First, the heights of the Longevity individuals were regressed (using [88]) against age and sex. Second, the residuals from the first step were regressed against their PS (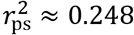, comparable to [12]; Supplementary Figure 2). The regression line from the second step was used to predict the height of the simulated offspring.

To generate the optimal PS, we first determined whether imputation had an effect on prediction accuracy. We used IMPUTE2 [89] and The Ashkenazi Genome Consortium reference panel [41]. Imputed data was post-processed to include only single nucleotide variants present in the summary statistics and with IMPUTE2 INFO-score >0.9. The 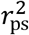 for height prediction (using all SNPs) was 0.201, which was slightly lower than for the PS generated without imputation, consistent with previous reports [90]. Since imputation incurs a significant computational and storage burden, we proceeded with the genotyped SNPs only.

Next, we considered the effect of linkage-disequilibrium (LD) pruning and P-value thresholds. LD-clumping was performed in PLINK [87] with window size of 250kb and *r*^2^ threshold of 0.1. LD was estimated based on 574 genomes from The Ashkenazi Genome Consortium [41], reduced to the 657,179 SNPs intersecting with the Longevity study. The number of remaining SNPs after LD-clumping was 93,345. We considered P-value thresholds between 10^−7^ to 1 in multiples of 10. We then searched for the parameter combination giving the maximum 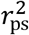 between predicted and actual phenotypes. Without LD-pruning, the maximal 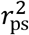 was 0.207 (using a P-value cutoff of 0.1). With LD-pruning, the maximal 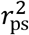 was 0.248, using a P-value cutoff of 0.001. Thus, our final score used LD-pruning and P<0.001, and included 15,752 SNPs.

#### General cognitive function

We used summary statistics from [11], based on a meta-analysis of intelligence (excluding the ASPIS cohort). Out of total of 9,145,263 SNPs, 468,809 intersected with the ASPIS panel. Following the results from height, we did not consider imputation. The optimal LD-clumping threshold and P-value threshold were *r*^2^ = 0.3 and 1, respectively, leaving 130,199 SNPs and reaching 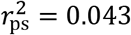 (Supplementary Figure 2). For improving the accuracy of LD estimation, we considered the entire 1066 genotyped individuals, including those without phenotypes.

We note that other approaches for genetic prediction may have slightly higher predictive power. However, an extensive benchmarking of methods and thresholds for trait prediction is beyond the scope of this paper. Our quantitative model allows us to approximate the utility of any score, based on its proportion of variance explained.

#### Simulating embryos

The Longevity cohort included actual couples, and these were used to simulate offspring (“actual matching”). For both the Longevity and the ASPIS cohorts, we also matched parents randomly (“random matching”). Given a pair of parents, we simulated offspring (embryos) by specifying the locations of crossovers in each parent. Recombination was modeled as a Poisson process, with distances measured in cM using the HapMap genetic map. For each parent, we drew the number of crossovers in each chromosome from a Poisson distribution with a mean equal to the chromosome length in Morgan. Random positions along the chromosome (in cM) represented the locations of the crossovers. We mixed the phased paternal and maternal chromosomes of the parent according to the crossovers’ locations, and randomly chose one of the resulting sequences as the chromosome transmitted from that parent. Note that due to phase switch errors, the paternal and maternal chromosomes are each a mixture of both. Nevertheless, phasing is expected to be accurate over short distances (switch error rate around 1%) [91], thus correctly representing LD blocks.

We repeated the process to generate either 10 or 50 embryos per couple (whether a true couple or randomly matched). The number of couples for random matches was such that the total number of embryos was 5000 (Table 1).

**Table 1.** A list of the sets of simulated embryos.

| Cohort | Phenotype | Matching | Number of matches | Number of offspring per couple |
| --- | --- | --- | --- | --- |
| Longevity | Height | Random | 500 | 10 |
| Longevity | Height | Random | 100 | 50 |
| Longevity | Height | Actual | 102 | 10 |
| Longevity | Height | Actual | 102 | 50 |
| ASPIS | Cognitive ability | Random | 500 | 10 |
| ASPIS | Cognitive ability | Random | 100 | 50 |

To calculate the polygenic scores for the synthetic embryos, we used the same summary statistics as for the parents. To predict the phenotype of the embryos, we used the regression model that we have generated from the parents. The predicted phenotype is already in its natural units (cm or IQ points). Adding sex- or age-specific mean values was unnecessary, as we considered only the differences between embryos attributed to their genetics.

#### Real nuclear families

We used 28 large nuclear Jewish families with an average of 9.6 adult offspring (full-siblings) per family who have completed their growth. The families were recruited in Israel and in the US after obtaining IRB approvals in both locations. Details on the cohort, measurements, and genotyping appear elsewhere [92]. In short, participants signed a consent form and filled a medical questionnaire (to ensure there were no medical conditions that could have affected their growth), and their heights were measured with four technical repeats at an accuracy of ±0.1cm. All 308 consented participants were genotyped on the Affymetrix Axiom Biobank array (≈630,000 SNPs). One from each of six pairs of monozygotic twins was excluded. Heights were corrected for age and age^2^, then standardized to *Z*-scores in each sex separately, then reported as 173.0 + 5.6*Z*cm.

For predicting height, we used the same set of 15,752 SNPs as used for the Longevity cohort, based on P<0.001 and LD *r*^2^ < 0.1. Of these, we used a total of 15,124 SNPs that were present on the array or could be imputed from the AJ reference panel [93]. We excluded SNPs homozygous in all participants. The weight of each SNP was its effect size [12], zero centered for the cohort, and the score of each subject was the weighted sum of the number of carried effect alleles. Scores were standardized into *Z*-scores and reported as for the actual heights.

## Acknowledgements

We thank Yaniv Erlich for discussions. S. C. thanks the Abisch-Frenkel Foundation for financial support. T. L. was supported, in part, by a grant from the National Institutes of Health (R01MH117646). The study of the nuclear families was supported by the James S. McDonnell Centennial Fellowship in Human Genetics to L. K.

## Supplementary Figures

**Supplementary Figure 1.**
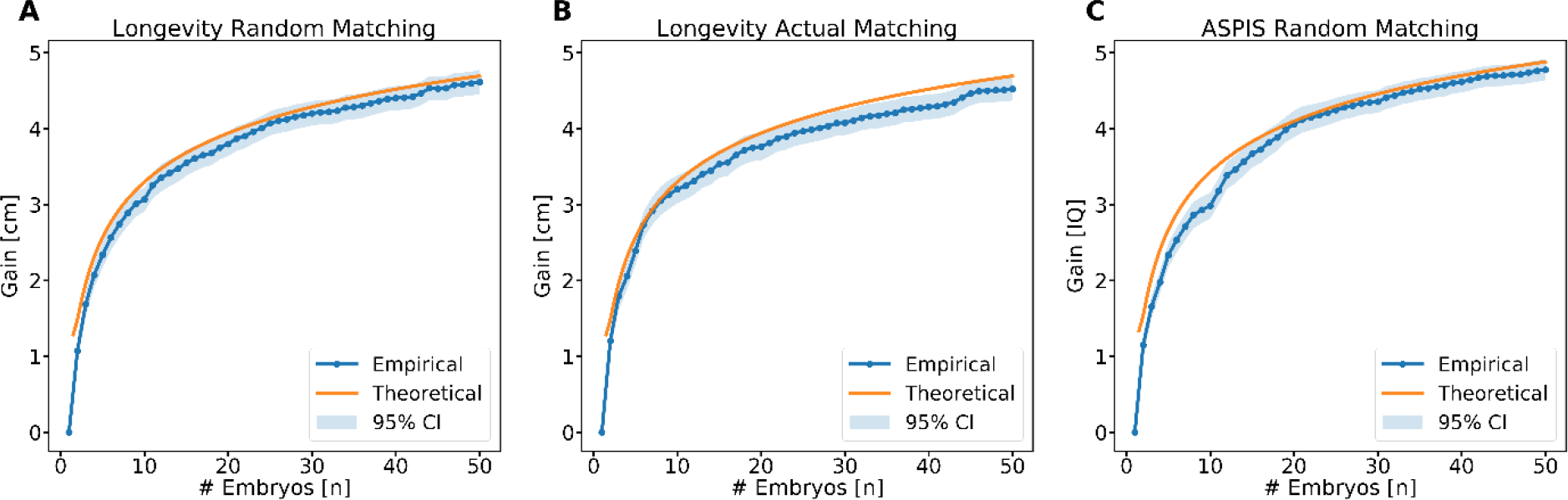
The mean gain in embryo selection vs the number of embryos *n*. All details are the same as in Figure 2. The theoretical prediction here is based on extreme value theory, as given in **Supplementary Note** Eq. (35), providing a slightly better fit compared to main text Eq. (1).

**Supplementary Figure 2.**
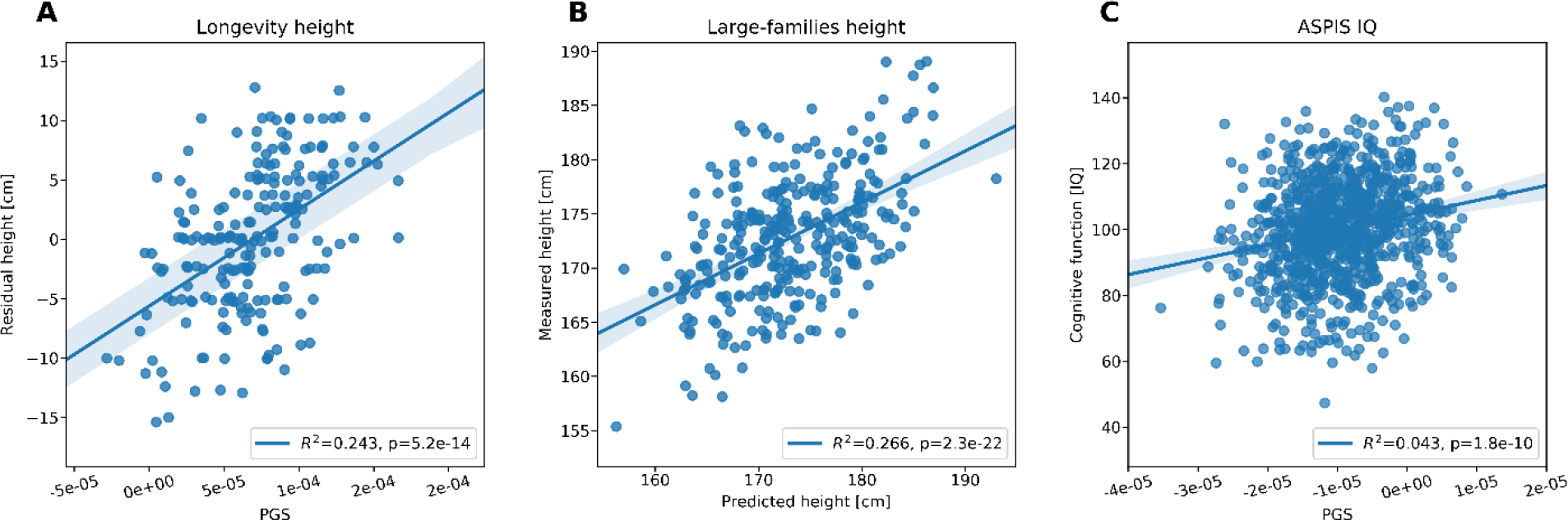
Height and cognitive ability (IQ) vs their polygenic scores. Results are shown for the heights of 204 individuals in the Longevity cohort **(A)**, the heights of 308 individuals from the large nuclear families **(B)**, and the IQ of 919 individuals from the ASPIS cohort **(C)**. Also shown are the regression lines, the proportions of variance explained, and the P-values. The proportions of variance explained by the polygenic scores are ≈25-27% for height and ≈4.3% for IQ.

## 1 Background and model

We assume a couple has generated *n* embryos, and we would like to select the optimal embryo with respect to a given polygenic trait. We assume that the genetic architecture of the trait is infinitesimal, namely that there are numerous causal variants, uniformly distributed along the genome. Denote the value of the trait as *z*, the number of variants as *N*, the variance of the trait as 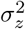, and the heritability as *h*^2^, and assume the trait has zero mean.

Mathematically, we assume an additive model, where for a given individual,

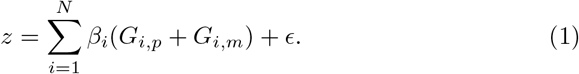

In the above equation, *G*_*i,p*_ = *g*_*i,p*_ − *f*_*i*_, where *g*_*i,p*_ ∈ {0, 1} is the number of minor alleles at site *i* on the paternal chromosome and *f*_*i*_ is the minor allele frequency. *G*_*i,m*_ is similarly defined for the maternal chromosome. *β*_*i*_ is the additive effect size per allele.

The polygenic score for the trait is defined as

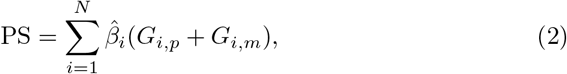

where the 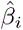s are the estimated effect sizes. We further assume that the trait can be modeled as

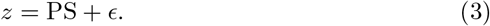

The error term now represents both the environmental component as well as unaccounted-for genetic components. The proportion of variance of *z* explained by the polygenic score PS is denoted

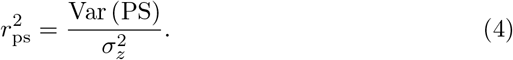

*r*_ps_ is also the correlation coefficient between the polygenic score and the trait value.

Next, we make the following assumptions. First, we assume that there is no assortative mating. This implies that *beyond linkage disequilibrium*, there is no correlation between the contributions to the polygenic score from *(i)* the two homologous chromosomes of an individual, at the same locus; *(ii)* two chromosomes of spouses, at the same locus; *(iii)* two distinct loci, coming from the same chromosome; and *(iv)* two distinct loci, coming from either two homologous chromosomes or from chromosomes of spouses. While assortative mating was demonstrated for several polygenic traits [1, 2, 3], our empirical data shows that the implied correlation between polygenic scores of spouses is relatively small. Specifically, we found that the correlation in the polygenic scores for IQ between actual spouses was relatively low and did not reach statistical significance (*r* = 0.12, *P* = 0.25). The correlation for the polygenic scores for height was similarly low (*r* = −0.03, *P* = 0.76). While the correlation may increase with the increasing predictive power of the scores, our model still serves as a useful baseline. In particular, since assortative mating is usually positive, our results form an upper bound for the utility of embryo selection.

Second, to avoid correlation due to linkage disequilibrium (LD), we write the polygenic score as a sum of *M* elements, where each element is the score in a single LD block,

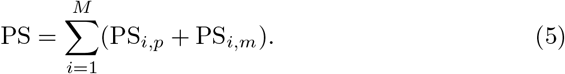

Above, 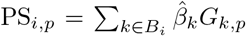, where *B*_*i*_ is the set of variants in block *i*, and similarly for PS_*i,m*_. Under the above assumption of no assortative mating, and assuming no correlation across LD blocks, this implies that for all *i* ≠ *j*, the random variables PS_*i,p*_, PS_*i,m*_, PS_*j,p*_, PS_*j,m*_ are all uncorrelated. Moreover, PS_*i,p,*_ PS_*i,m*_ for any one individual are uncorrelated with PS*i,p* and PS_*i,m*_ in the spouse of that individual, for any block *i*. The LD blocks can be identified, e.g., as in [4].

We further assume that all blocks contribute equally to the variance (although this can be easily relaxed, leading to the same result). Thus, under the above model, we have

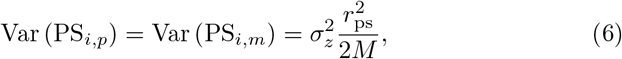

as well as

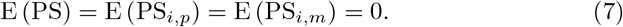

Next, we consider the vector PS = (PS^1^, …, PS^*n*^) of polygenic scores for *n* embryos together. We assume that the distribution of the polygenic scores, PS, is normal in each embryo (due to the polygenic nature of most complex traits [5]), and further that the joint distribution of the polygenic scores over *n* embryos is multivariate normal,

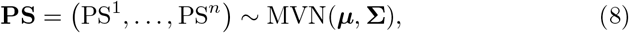

where ***μ*** = **0**_*n*_ (a column vector of zeros of length *n*). The diagonal elements of the covariance matrix **Σ** are 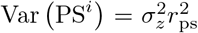 for all *i* = 1, …, *n*. We will compute the off-diagonal covariances below (Section 2).

We define the *gain G* due to embryo selection as the difference between the polygenic score of the best embryo and the average scores of all embryos. Mathematically,

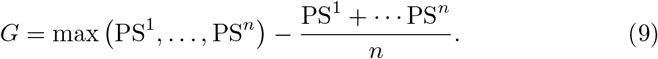

The gain *G* is a random variable, with a sample space over all theoretical sets of *n* siblings. In the following, we will examine the statistical properties (e.g., mean and variance) of the gain as a function of *n*, 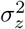, and 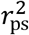.

For the mean gain, using Eq. (7),

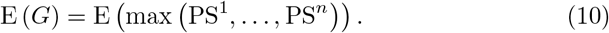

We derive an approximate formula for the mean gain in Section 3. We then consider in Section 4 other properties of the gain *G*, including its variance and the implications for prediction of the embryo with the actual highest trait value.

## 2 The covariance

In order to obtain the joint distribution of (PS^1^, …, PS^*n*^), we need to compute Cov (PS^*A*^, PS^*B*^), the covariance between the polygenic scores of two distinct embryos (or siblings), which we name *A* and *B*. For two individuals *A, B* with kinship coefficient Θ, standard quantitative genetics theory gives the covariance Cov (*z*_A_, *z*_B_) = 2Θ*h*^2^, for a quantitative additive trait *z* with heritability *h*^2^ under the infinitesimal model [6]. Specifically, for full siblings, Θ = 1/4, and thus Cov (*z*_A_, *z*_B_) = *h*^2^/2. For completeness, we derive the corresponding result here for the polygenic scores PS^*A*^ and PS^*B*^.

Recall that we modeled the polygenic score as 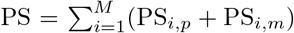, where PS_*i,p*_ is the score of the *i*^th^ LD block in the paternal chromosome and PS_*i,m*_ is the score from the maternal chromosome. For a pair of siblings and for a given LD block, their scores come from the same parental chromosome with probability 1/2, or from different parental chromosome with probability 1/2. (We ignore the possibility of a recombination event taking place in the middle of an LD block, because, first, by definition, recombination is depleted within LD blocks, and second, the distance between crossovers is much greater than the distance between LD blocks [7].)

Consider the two homologous chromosomes of the father at block *i*. Denote the polygenic score of the first chromosome (say, grandpaternal) as *x*_*i*,1_ and the score of the second chromosome (say, grandmaternal) as *x*_*i*,2_. Similarly, denote the polygenic scores of the two maternal chromosomes as *y*_*i*,1_ and *y*_*i*,2_. For embryo *A*, denote by *p*_*A,i*_ the choice of the paternal chromosome transmitted to embryo *A* at block *i*: *p*_*A,i*_ = 1, 2 with equal probability. Similarly, *m*_*A,i*_ = 1, 2 denotes the identity of the maternal chromosome transmitted to embryo *A* at block *i*. With the above notation, the polygenic score of embryo *A* can be written as:

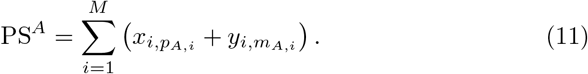

Similarly,

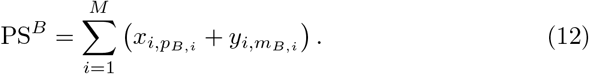

The covariance between the scores of two embryos is

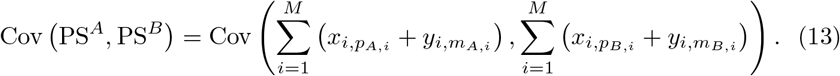

According to the assumptions of Section 1, there is no correlation between the scores of any two blocks on two chromosomes of spouses, or between distinct blocks on the same chromosome. Thus,

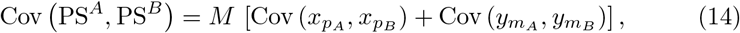

where *p*_*A*_, *p*_*B*_, *m*_*A*_, *m*_*B*_ are the identities of the chromosome transmitted by the father/mother to embryos *A* and *B* at a representative block, and *x*_1_, *x*_2_, *y*_1_, *y*_2_ are the scores of the four parental chromosomes in that block. *p*_*A*_, *p*_*B*_, *m*_*A*_, *m*_*B*_ are independent random variables taking the values 1 or 2 with equal probabilities. To compute the remaining terms, we invoke the law of total covariance, by conditioning on *p*_*A*_, *p*_*B*_ or on *m*_*A*_, *m*_*B*_. For example,

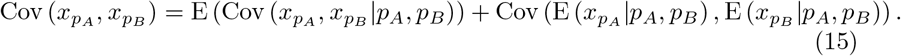

However, E (*x*_*pA*_|*p*_*A*_, *p*_*B*_) = E (*x*_*pB*_|*p*_*A*_, *p*_*B*_) = 0, and are both in general independent of *p*_*A*_ or *p*_*B*_. Thus, the second term (covariance of expectations) vanishes. We can expand the first term as follows,

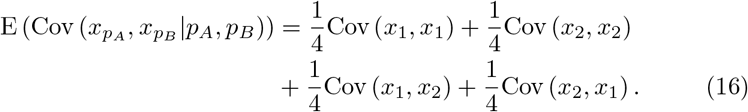

Again according to the assumptions of Section 1, there is no correlation between the scores of blocks from homologous chromosomes. Thus, the two terms in the second line vanish. Finally, using Eq. (6),

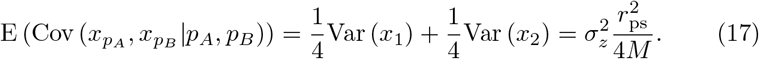

A similar result holds for the maternal scores. Using Eq. (14),

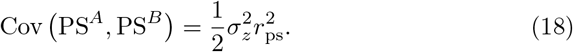

We have thus specified the distribution of the polygenic scores of the *n* embryos,

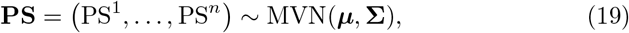

where ***μ*** = **0**_*n*_ and **Σ** is an *n* × *n* covariance matrix with elements

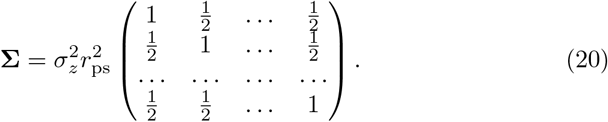

## 3 The mean score of the top-scoring embryo

Define PS_max_ = max (PS^1^, …, PS^*n*^). The mean gain (as defined in Section 1) is the mean of the score of the top-scoring embryo, E(*G*) = E (PS_max_) (Eq. (10)).

Written more generally, we would like to compute the mean of the maximum of *n* multivariate normal variables, denoted **PS** = (PS^1^, …, PS^*n*^) ~ MVN(**0**_*n*_, **Σ**), where the covariance matrix **Σ** is defined according to Eq. (20). We can write the covariance matrix also as **Σ** = ***A*** + ***B***, where

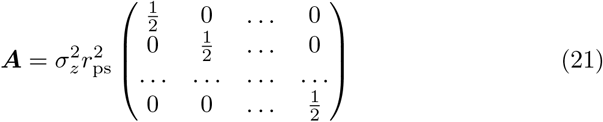

and

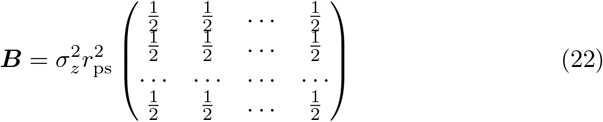

Given this decomposition, we can write the distribution of polygenic scores as a sum of two independent multivariate normal variables **PS** = ***Y*** + ***Z***, where

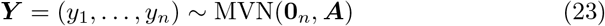

and

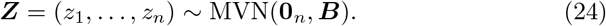

The covariance matrix ***A*** of ***Y*** is diagonal, and hence the variables in ***Y*** are independent. ***Z*** has a constant covariance matrix ***B***, which means that the correlation between all variables is 1. Thus, all elements of ***Z*** are equal to the same normal variable with variance given by Eq. (18),

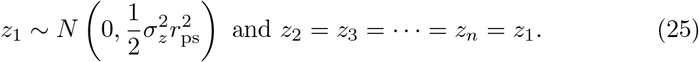

Since **PS** = *Y* + *Z*, we have

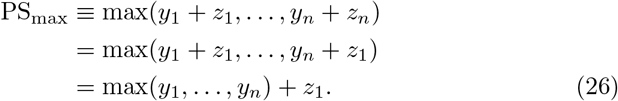

The expectation of PS_max_ is

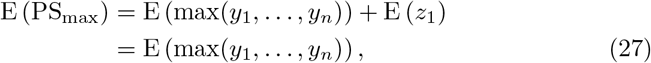

since *Z* has zero means. Therefore, the mean of the maximum of (PS^1^, …, PS^*n*^) is the same as the mean of the maximum of *n independent* normal variables with variance 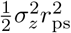 each.

For independent normal variables, some results are known for the expectation of the maximum. For example, in [8] it was shown that if *R* = max(*x*_1_, …, *x*_*n*_), where *x*_*i*_ ~ *N*(0, *σ*^2^) are independent, then

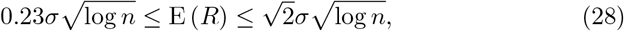

and thus for very large *n*,

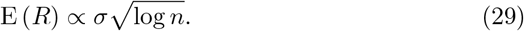

Numerically, we found the best fit to Eq. (29) (over *n* from 1 to 50) was when the coefficient of proportion was ≈ 1.09. In our case, 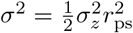. Noting that 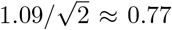, the mean polygenic score of the best embryo, and hence the mean gain, is

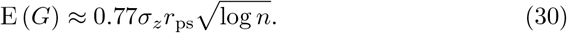

Due to its simple functional form, we report Eq. (30) as Eq. (1) of the main text. However, these bounds are not tight. Based on extreme value theory, we can reach a more accurate expression. For large *n* [9, 10], the maximum of *n* standard normal variables has an approximate *Gumbel* distribution with CDF:

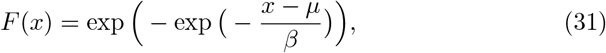

where

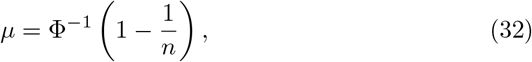

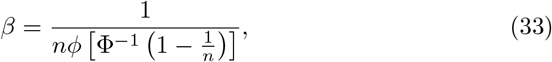

*ϕ* is the PDF of the standard normal distribution, and Φ^*−*1^ is the inverse CDF of the standard normal distribution. The mean of a Gumbel random variable is *μ* + *βγ*, where *γ* is the Euler-Mascheroni constant (*γ* ≈ 0.577).

In our case, all normal variables have standard deviation *σ*, and thus,

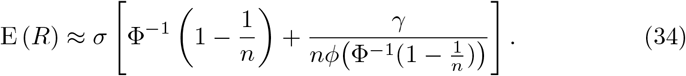

Finally, as we have 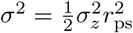,

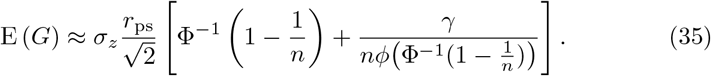

We found this equation to be more accurate than Eq. (30) (Supplementary Figure 1).

## 4 Additional calculations

### 4.1 The variance of the score of the top-scoring embryo

Extreme value theory can also provide an expression for the variance of the top score. From Eq. (26),

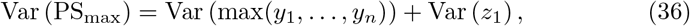

where 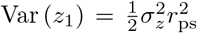. The variance of a Gumbel variable is known to be *π*^2^*β*^2^/6. Thus,

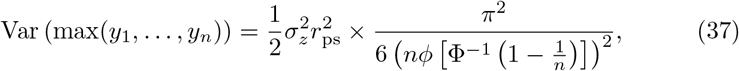

and

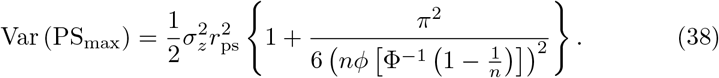

Eq. (38) is the variance of the best polygenic scores among the *n* embryos. However, it does not provide us the variance of the gain *G*. To compute the variance of the gain, we would need to compute the covariance between the maximum score and the other scores, which we leave to future work.

### 4.2 A prediction interval for the phenotype

We have so far predicted the mean value of the score of the top-scoring embryo (Eq. (30)). However, even for a given polygenic score of the best embryo, the actual value of the trait may differ considerably. Denote the value of the trait of the top-scoring embryo as

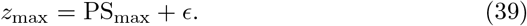

Following Section 1, *ϵ* has zero mean and variance

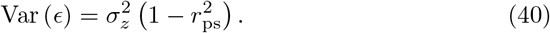

Thus, given its polygenic score PS_max_, the remaining variance in trait value for the top-scoring embryo is 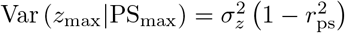. Assuming a normal distribution for *ϵ*, a 95% prediction interval for the actual value of the trait will be approximately

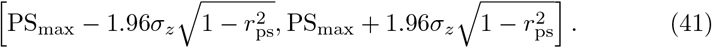

Eq. (41) is Eq. (2) in the main text. The above prediction interval is centered around PS_max_, which is assumed to be known. When it is unknown, a reasonable approximation for the center of the prediction interval may be *z*_mp_+*E*(*G*), where *z*_mp_ is the mid-parental trait value (i.e., the average of the (sex-adjusted) trait between the two parents). Theoretically, this approximation should break down for the most extreme tails of parental phenotypes, because the gain must be smaller in these cases. However, our simulations (Supplementary Note Figure 1) suggest that in a realistic setting, the gain does not significantly depend on the mid-parental trait value.

In a naïve calculation for no selection, we assume no information is available regarding the embryo, and thus, the 95% prediction interval would be

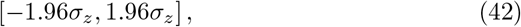

as for any normal variable with zero mean and variance 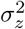. However, the phenotype can be predicted based on the mid-parental trait value. Denote the trait of an offspring as *z*_*o*_. A well-known result in quantitative genetics is that the slope of the regression of *z*_*o*_ on *z*_mp_ is equal to the heritability *h*^2^ [6]. The correlation coefficient is the product of the slope and the ratio of the standard deviations, 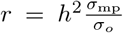. But 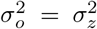 and 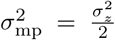. Thus, 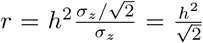. The proportion of variance explained is 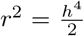 (see also, e.g., [11]), and the remaining variance is 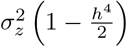. Thus, a more realistic 95% prediction interval for the case of no selection would be

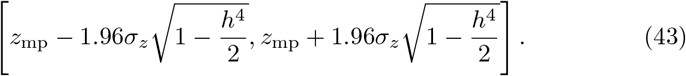

In theory, having both the mid-parental value and the offspring’s PGS may lead to a more accurate prediction, with a narrower prediction interval, even for the case of selection. Prediction in this setting is in general non-trivial, and more so here since the embryo is non-random but rather selected for its high polygenic score. The combination of both the polygenic score and the mid-parental value cannot explain more variance than implicated by the heritability. Thus, the proportion of variance explained by all of the available data (PS and parents’ trait value) can be anything within the range 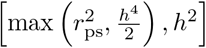, i.e., it is at least the best of the two predictors, but no higher than the heritability.

**Figure 1:**
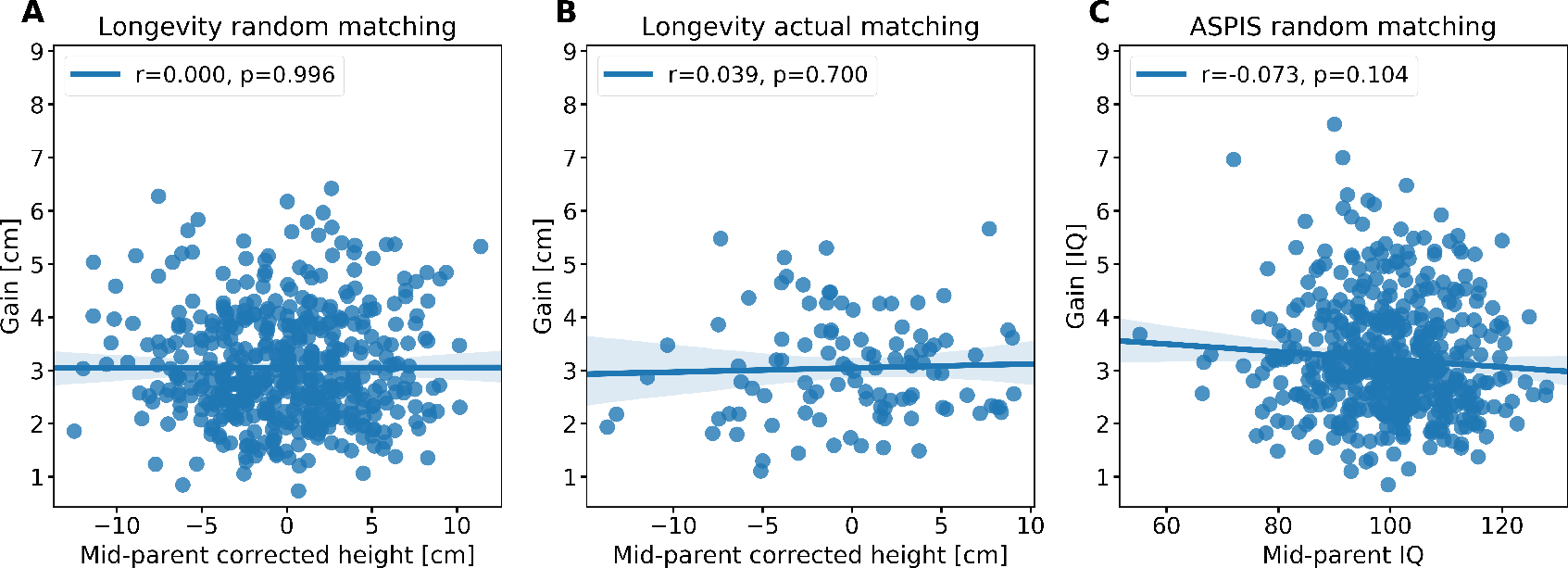
The gain in embryo selection vs the mid-parental trait value. The height was corrected for sex and age. The height residuals or the IQ points were then averaged between the two parents. (A) Random mating for height. (B) Actual couples for height. (C) Random mating for IQ. The gain was calculated over *n* = 10 embryos. The correlation coefficient and its associated P-value are shown at the top of each panel.

At present, the variance explained by the mid-parental trait value is about the same as that explained by the PS for height, but much higher than that explained by the PS for many other traits, including cognitive ability. In the future, the variance explained by the PS may substantially exceed that explained by the parents. In our main text examples, we consider predictors explaining 70% of the variance in height and 30% of the variance in cognitive ability — these are much larger proportions compared to those explained by the mid-parental height or IQ: 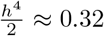 for height (assuming *h*^2^ ≈ 0.8) and 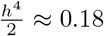 for IQ (assuming *h*^2^ ≈ 0.6). In these extreme cases, the prediction interval in Eq. (41) probably cannot be made substantially narrower.

### 4.3 The mean difference between the top-ranked trait and the trait of the best embryo

In the main text, we analyzed real large nuclear families. When reduced to *n* = 7 children per family, we found that the average height difference between the tallest child and the child with the best PS was 3.0cm. To determine the expectation based on our quantitative model, consider *n* siblings, whose PSs are modeled as a multivariate normal variable,

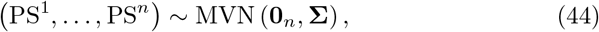

where **Σ** is defined in Eq. (20). We assume that the phenotypes, *z*_1_, …, *z*_*n*_, can be modeled as

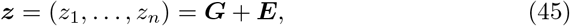

where ***G*** ~ MVN (**0**_*n*_, **Σ**_*g*_) and ***E*** ~ MVN (**0**_*n*_, **Σ**_*e*_), with

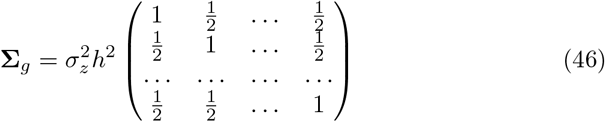

and

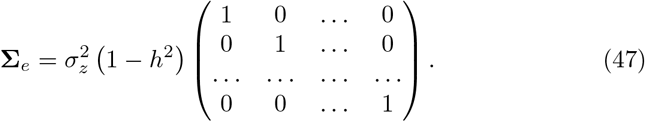

In the matrix **Σ**_*g*_, the off-diagonal elements are 1/2 due to the covariance between sibs, as in Section 2. We assume no covariance between the environmental components. Thus, in total, (*z*_1_, …, *z*_*n*_) ~ MVN (**0**_*n*_, **Σ**_*z*_), where

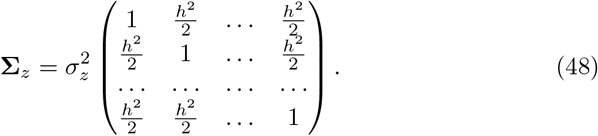

As we have shown in Section 3, because the covariance terms are all equal, the mean of the maximum of the phenotypes (***z***) is equal to the mean of the maximum of *n* independent normal variables, each with zero mean and variance 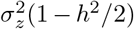. Denote by E(*R*) the mean of the maximum of *n* standard normal variables (e.g., as we calculated in Eq. (34)). Denote the maximum phenotype across the sibs as *z*_*m*_. Since the identity of this sib is not known at the time of selection, the phenotype of the selected embryo, *z*_max_, may be lower, and we have (using Eq. (39)),

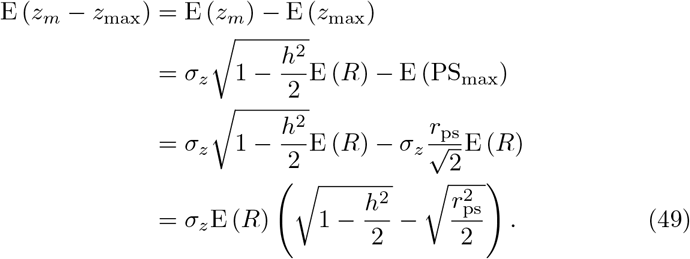

We obtained E (*R*) exactly based on numerical integration, substituted *σ*_*z*_ = 5.6cm, *h*^2^ = 0.8, and 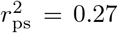, and obtained E (*z*_*m*_ − *z*_max_) = 3.1cm, very similar to the observed value.

### 4.4 The probability of the top-ranked embryo to have the top-ranked trait

When reduced to *n* = 7 children per family, we found in the real data that on average, in ≈ 31.5% of the families the child whose PS was ranked first was also ranked first in actual height. To determine the expectation based on our quantitative model, consider again *n* siblings. Recall that their phenotypes, ***z*** = (*z*_1_, …, *z*_*n*_), are modeled as

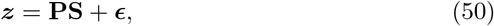

as in Eq. (39). The polygenic scores are as defined above (Eq. (44)). For the error term, we have ***ϵ*** ~ MVN (**0**_*n*_, **Σ**_*ϵ*_), and

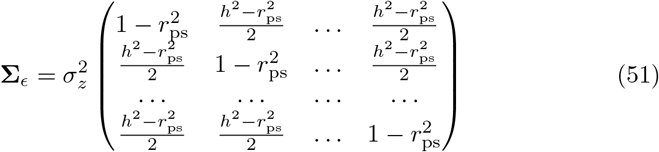

To explain the above equation, each *ϵ*_*i*_ has variance 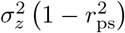. However, here the *ϵ*_*i*_’s must be correlated because they model not only the environment but also the genetic component not modeled by the PGS. The off-diagonal entries in the covariance matrix of the phenotypes ***z*** are equal to 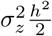 from Eq. (48). Assuming independence between **PS** and ***ϵ***, these entries are equal to the sum of the off-diagonal entries in the covariance matrix of **PS**, 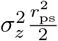 (Eq. (18)), and the off-diagonal entries in the covariance matrix of ***ϵ***. Thus, the latter must be 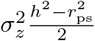.

We simulated values for **PS** and ***ϵ***, assuming *n* = 7, *h*^2^ = 0.8, and 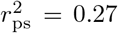 as in the real family data, and then calculated the phenotypes according to Eq. (50). (The value of *σ*_*z*_ does not change the relative ranks, and can be set to any value.) We found that in ≈ 33.4% of the simulations, the sibling top-ranked for the score (PS) was also top-ranked for the phenotype (*z*), in a reasonable agreement with the empirical results. An analytic approximation to this probability can also be derived based on Eq. (14) in [12].

